# Ribose lowers RpoS translation through RbsD mRNA

**DOI:** 10.1101/2023.08.28.555190

**Authors:** Andrew Badaoui, Isabel Smith, Melisa Balla, Mikayla Cavanaugh, Julia Lockart, Dylan Parsons, Esteban Martes, Shaina Dorean, Cheik Diop, Thomas Tran, Celeste N. Peterson

**Author notes:** Corresponding Author: Celeste N. Peterson. First authors IS and AB contributed equally.

## Abstract

The *Escherichia coli* sigma factor RpoS accumulates during starvation, activates transcription of the general stress response, and then rapidly returns to low levels upon nutrient replenishment. Sugar levels modulate RpoS levels by signaling though central metabolism, ATP and the phosphoenolpyruvate-carbohydrate phosphotransferase system (PTS) transport system, which favors glucose. However, there have been few examples of dedicated control pathways for specific sugars regulating RpoS. We screened an overexpression library with a RpoS’-‘LacZ reporter and found that overexpressing the ribose metabolism *rbsD* gene reduced RpoS levels. A functional RbsD ribose pyranase protein was not necessary for the effect, suggesting that the RbsD mRNA was responsible for the effect. We used a series of LacZ fusions and RT-qPCR to determine how RbsD mRNA affects RpoS and found that regulation of RpoS occurred at the level of translation. Furthermore, the effect of *rbsD* overexpression was diminished in strain that did not have the RpoS untranslated hairpin loop or the small RNA chaperone Hfq. RbsD mRNA has previously been shown to bind to the sRNA DsrA. We demonstrated that the effect of RbsD on RpoS was dependent on the sRNAs DsrA and ArcZ and Hfq. Finally, we showed that the sugar ribose lowers RpoS levels in a manner that requires RbsD. In summary, our results demonstrate that RpoS levels respond specifically to ribose though a component of the ribose metabolism pathway and the main player, RbsD, acts as dual function mRNA that has a regulatory role by interacting with the sRNAs that control RpoS.

## Importance

Bacteria live in diverse environments and need to respond quickly to fluctuating carbon availability. In Escherichia coli, the sigma factor responsible for survival during starvation is RpoS. For sugars, both the general carbohydrate transport process and central metabolism components regulate RpoS. However, despite the importance of RpoS to the carbon response, it is still unclear if different sugars have different regulatory pathways. We show that RpoS levels respond to ribose through a specific regulatory pathway that does not involve central metabolism. Specifically, one of the components of ribose metabolism has a dual purpose whereby it converts ribose to a different form and separately downregulates RpoS translation. This dedicated pathway highlights ribose as an important signal for exit from stationary phase.

## Introduction

Exposure to one stress, such as starvation, leads to cross resistance against a wide variety of other stresses in *Escherichia coli*. Upregulation of this general stress response can not only infer protection against disparate stressors such as carbon starvation, oxidative stress, osmolarity stress but can also result in increased tolerance to antibiotics (1, 2). The master regulator of the general stress response is the alternative sigma factor RpoS. By binding to the core RNA polymerase and contacting the -10 and -35 promoter sequences, RpoS controls transcription of hundreds of genes (3). One of the main stressors inducing RpoS is carbon starvation, and upon carbon replenishment, RpoS levels rapidly decrease. To date, signaling molecules regulating RpoS in response to fluctuating carbon levels have been linked to central metabolism, primarily through ATP, actetyl-CoA and the alarmone ppGpp (4–6). Glucose affects RpoS expression via the phosphoenolpyruvate-carbohydrate phosphotransferase system (PTS), which prefers glucose and affects ppGpp production, and via cAMP, but there are conflicting reports on whether cAMP activates or represses RpoS transcription (4, 7). Dedicated regulatory pathways connecting RpoS to specific sugar levels have seemed strikingly absent.

Regulation of RpoS primarily occurs at the post-transcriptional level, which allows for a rapid response to stress. RpoS translation is positively controlled by three known sRNA interacting with sRNA chaperone Hfq – DsrA, ArcZ and RprA – in response to e.g. cold temperature, membrane stress or anaerobic conditions (2, 8). These sRNAs work by increasing accessibility to the 5’ UTR of RpoS so that the ribosome can bind and initiate translation. Without the sRNAs, the 5’UTR forms a hairpin loop that occludes the ribosome binding site. The sRNAs also help stabilize the mRNA and prevent premature transcriptional termination (9, 10). There is also RpoS regulation independently of the sRNAs; for example, acetyl-CoA regulates RpoS translation in a manner that does not require any of the three known sRNAs (4). RpoS is also regulated at the level of protein degradation. When conditions are plentiful, RpoS is made but targeted for degradation by an adaptor protein SprE (RssB) and the protease ClpXP. However, during starvation, RpoS degradation ceases and the protein accumulates. (2, 11). There are several anti-adaptor proteins that affect SprE (RssB) activity and the degradation process itself is affected by ATP and acetyl-CoA levels (4, 5, 12, 13).

All three RpoS-regulating sRNAs – DsrA, ArcZ and RprA – have multiple targets in addition to RpoS. DsrA is a 90 base pair RNA and its first stem loop forms an RNA-RNA complex with the RpoS UTR that allows for RpoS translation (14). The DsrA structure is dynamic and is also capable of binding to other targets including *rbsD*, *hns*, *mreB* in *E. coli* and the *pflB* metabolism gene in *Salmonella* Typhimurium. (15–17). The *rbsD* gene codes for the RbsD protein that works in concert with RbsK for the conversion of d-ribose into d-ribose 5-phosphate (18). DsrA binding to RbsD causes degradation of the RbsD mRNA (16). Notably, DsrA binds to a region approximately three quarters along the *rbsD* gene, far from the start codon (16, 17). ArcZ also has additional targets including MutS and FlhDC (19, 20). Similarly, RprA also regulates several genes involved in biofilm formation and acid resistance (21). Thus, these sRNAs form part of a complex network that allows the cell to integrate different environmental signals into multiple physiological outcomes.

While many different carbon sources can affect central metabolism and thus the exit from stationary phase, there is little evidence for pathways specific for a particular carbon source. To better identify the pathways involved in exit from starvation, we used an overexpression library to identify down-regulators of RpoS750’-‘LacZ. We identified RbsD as a negative regulator of RpoS translation. The RbsD mRNA, rather than the protein, regulated RpoS through the sRNAs DsrA and ArcZ. Finally, ribose lowered RpoS expression in a manner that required RbsD.

## RESULTS

### Overexpression of *rbsD* mRNA lowers RpoS levels

In order to identify novel regulators of RpoS, we carried out an overexpression screen with an LacZ reporter for RpoS (22). Briefly, a library was made by digesting the *E. coli* genome with *EcoRV,* purifying fragments that were between 500 and 1000 bases and cloning them into a low copy vector. This library was transformed into an RpoS750’-‘LacZ strain (22), which reports RpoS transcription, translation, and degradation. Approximately 30,000 colonies were screened for changes in color on Xgal/LB/kan plates and one lighter blue colony was selected for further study. The plasmid was re-transformed into the original reporter strain, and it was confirmed that the plasmid was the cause of the light blue phenotype rather than a chromosomal mutation. Sequencing revealed that the plasmid in this colony contained the full-length ribose pyranase gene *rbsD* gene and small parts of other neighboring genes, *tkrD* and *rbsA*. Subsequent cloning showed that the overexpression of the *rbsD* gene alone was sufficient for the phenotype.

We next measured the effect of *rbsD* on RpoS through betagalactosidase assays and Western blots. Overexpression of *rbsD* resulted in significantly lower RpoS expression than the wild type RpoS750’-‘LacZ strain in rich media (Figure 1a). To confirm that RpoS protein levels were indeed lowered, we followed up with a Western blot. Figure 1b shows that RpoS protein levels were also lowered with the *rbsD* overexpression plasmid. Thus, *rbsD* mRNA overexpression leads to lower RpoS protein levels.

**Figure 1.**
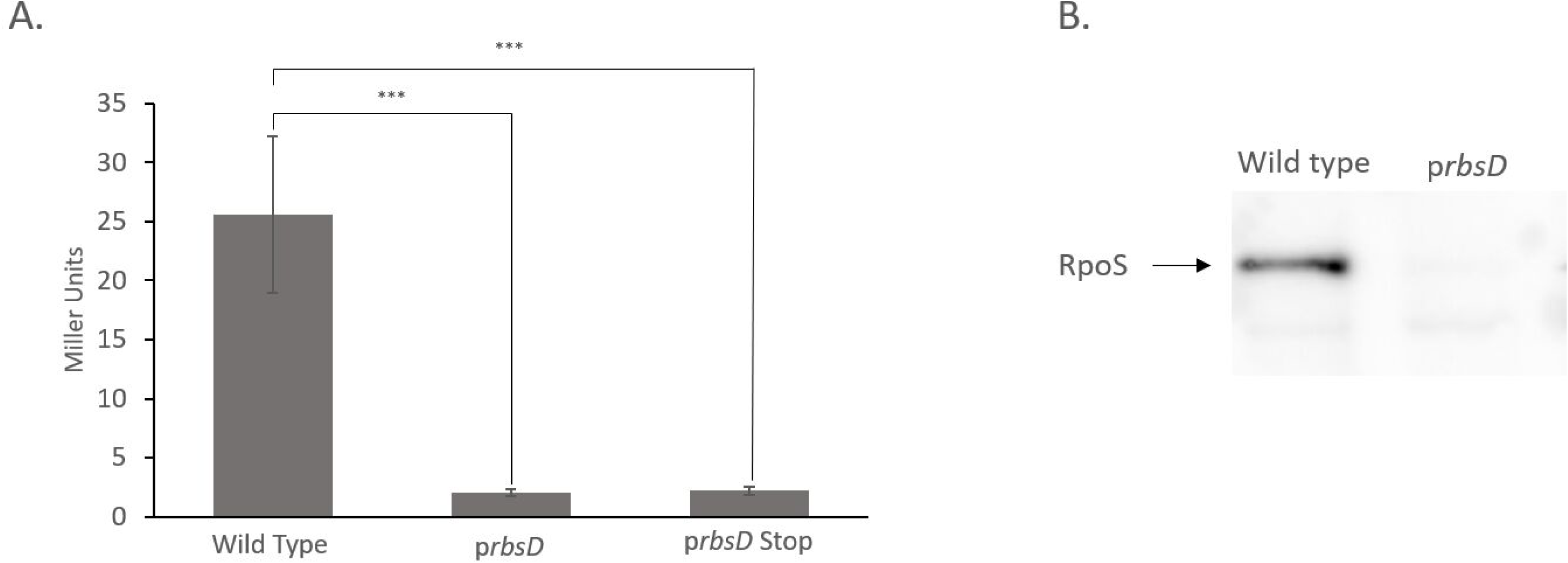
Overexpression of RbsD mRNA lowers RpoS levels. (A). Beta-galactosidase assays of the RpoS750’-‘LacZ fusion with an empty vector were compared to fusion strains carrying p*rbsD* and p*rbsDstop* plasmids. Exponentially growing cell in LB rich media were collected for the beta-galactosidase assays. Data show is an average of three independent experiments (***, P < 0.001). (B) Western blot of the wild type and the p*rbsD* stains growing exponentially in LB. Samples were preparation as described in Materials and methods section.

The RbsD mRNA codes for a ribose metabolism protein and also binds to the sRNA DsrA and the sRNA chaperone Hfq. To determine which RbsD function was responsible for the effect on RpoS, we introduced a stop codon in the *rbsD* gene on the overexpression plasmid. When the GGA codon (at position 96) was replaced with a UAA stop codon, the p*rbsDstop* plasmid had the same effect as p*rbsD* on RpoS’-‘LacZ levels (Figure 1a). This demonstrated that the protein coding part of RbsD mRNA is not necessary for lowering RpoS levels, and suggests that mRNA of DsrA plays a role.

### *rbsD* overexpression lowers RpoS translation in a manner that requires Hfq and the hairpin loop

To determine if the effect of RbsD on RpoS was occurring at the level of transcription, translation or protein stability, we used a series of lacZ fusions. The *prbsD* plasmid was identified with the long RpoS750’-‘LacZ fusion, which reports all three levels of control and is synthesized and degraded like the wild type RpoS (22). A second fusion, the RpoS477’-‘lacZ protein fusion, lacks the turnover element K143, so the reporter is not subject to RpoS degradation control (23). The p*rbsD* lowered levels of fusion to a similar fashion as the long fusion, indicating the RbsD is not regulating RpoS degradation. We the examined a transcriptional fusion, RpoS477’-LacZ+ (22). We found that there was fusion was not significantly downregulated with p*rbsD* (figure 2a). There was also not a very big effect on regulation as measured by RT-PCR, which showed RpoS mRNA were 2.2-fold lower with *prbsD* than the empty vector control. The inhibition of translation is often associated with a decrease in mRNA levels, potentially due to the mRNA being more susceptible to degradation without the ribosomes (9). Overall, our results are consistent with *rbsD* overexpression controlling RpoS translation, with a possibly indirect effect on mRNA levels.

**Figure 2.**
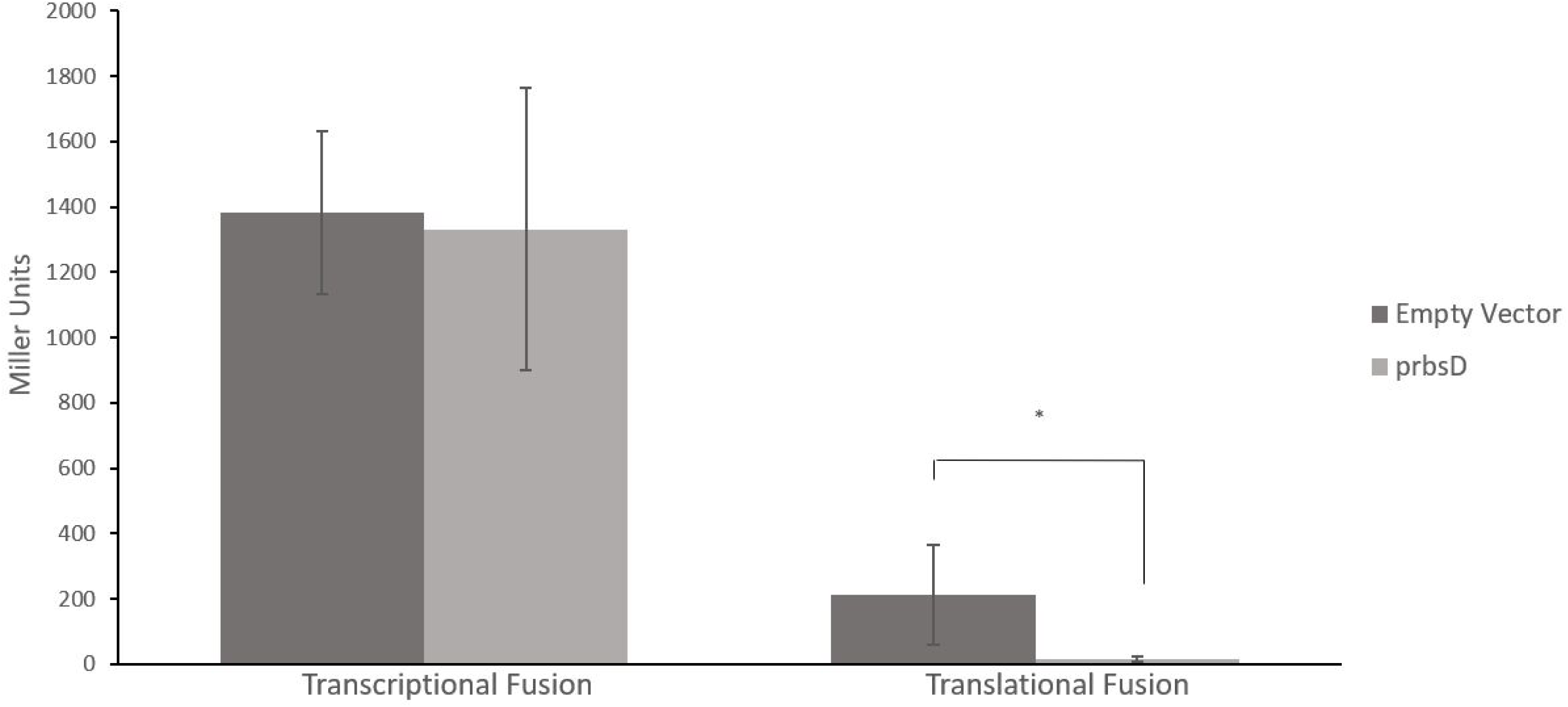
Overexpression of RbsD mRNA lowers RpoS levels in a post-transcriptional manner. (A) Beta-galactosidase assays with the RpoS-‘Lac+ transcription fusion showed no significant difference with and without the p*rbsD* overexpression plasmid. (B) Beta-galactosidase assays with RpoS477’-‘LacZ translation fusion, which reports both transcription and translation of RpoS but not degradation, showed diminished activity with the p*rbsD* overexpression plasmid (*, P < 0.05).

To confirm that RpoS translation was being regulated by p*rbsD* and to probe the role of the UTR hairpin turn, we used a pCP17-RpoS’-LacZ fusion. This fusion only reports RpoS regulation of translation through the UTR hairpin turn; it has neither the RpoS promoter nor the RpoS degradation element (4). Again, overexpression of *rbsD* lowered the levels of this fusion significantly, confirming regulation is at the level of translation (Figure 3a). A variant of this version has been created that includes a C125T mutant in the leader portion, disrupting the hairpin loop and regulation by Hfq and the sRNAs (4). The plasmid overexpressing *rbsD* did not significantly affect this mutated fusion compared to wild type (Figure 3a), suggesting that regulation by RbsD required the RpoS hairpin loop. In addition, an *hfq::cam* allele also abolished the effect of *rbsD* overexpression on the long RpoS750‘-‘LacZ fusion (Figure 3b). Thus, RbsD overexpression regulated RpoS translation through the hairpin loop and Hfq, suggesting that a sRNA might be involved.

**Figure 3.**
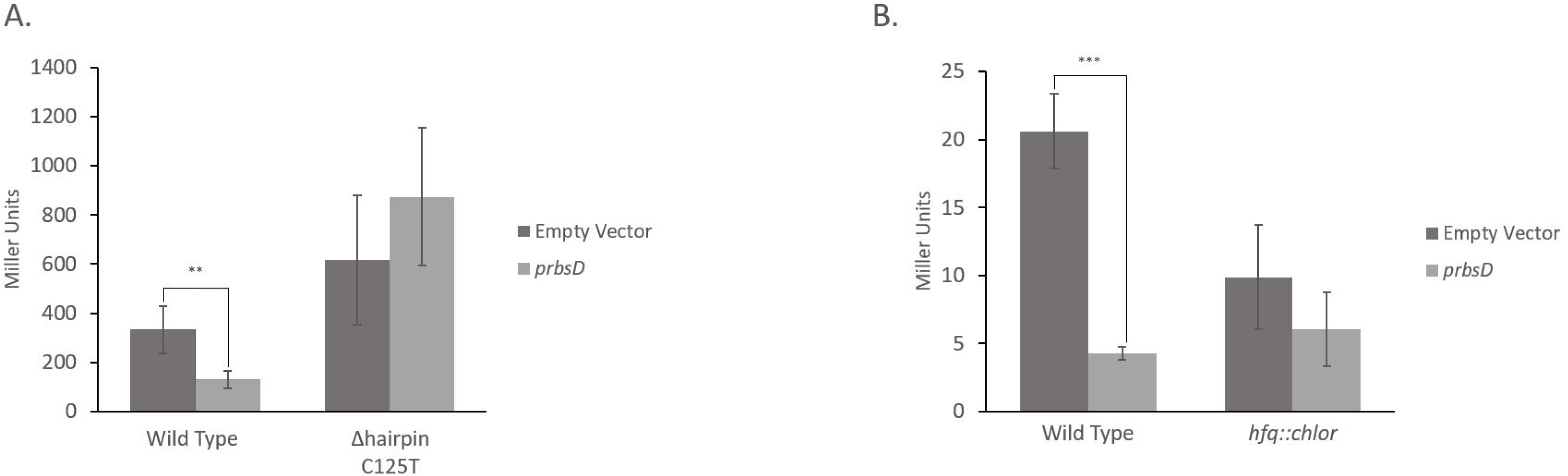
RbsD requires *rpos* hairpin loop and Hfq for translation control. (A) Beta-galactosidase assays with the pCP17-RpoS477’-‘LacZ fusion under the constitutive CP17 promoter, which reports only RpoS translation. This translation reporter showed lower activity with the p*rbsD* overexpression plasmid (**, P < 0.01; ***, P < 0.001). A derivative of the above reporter, Cp17-C125T mutant-rpoS477-lacZ, has the C125T mutation which disrupts the *rpoS* hairpin loop. This mutated reporter does not show any significant effect of the p*rbsD* overexpression plasmid. (B) Beta-galactosidase assays with the Rpos750’-‘LacZ fusion in the presence and absence of *hfq::chlor*. When the *hfq::chlor* is present, then the p*rbsD* overexpression plasmid no longer lowers Rpos750’-‘LacZ levels.

### DsrA and ArcZ sRNAs are required for RbsD regulation of RpoS

Using a series of sRNA deletions, we examined which sRNAs are required for the effect of p*rbsD* on RpoS. We examined the three sRNAs that are known to have a positive significant effect on RpoS: DsrA, ArcZ and RprA. We examined single sRNA mutants and also combined double mutants. For these experiments, the *rbsD* gene was cloned on to a spec resistant plasmid that has a higher copy number than the original kan resistant plasmid, resulting in a larger downregulation of RpoS. Figure 4a shows the effect of the single and double mutants on the RpoS477’-‘LacZ fusion. In the absence of *dsrA*, the effect of overexpression of *rbsD* was reduced from ∼16 fold to ∼5 fold. In the absence of both *dsrA* and *arcZ,* the effect of *rbsD* overexpression was abolished. Thus DsrA, and to a lesser extent ArcZ, played major roles in the regulation of RpoS by RbsD mRNA.

**Figure 4.**
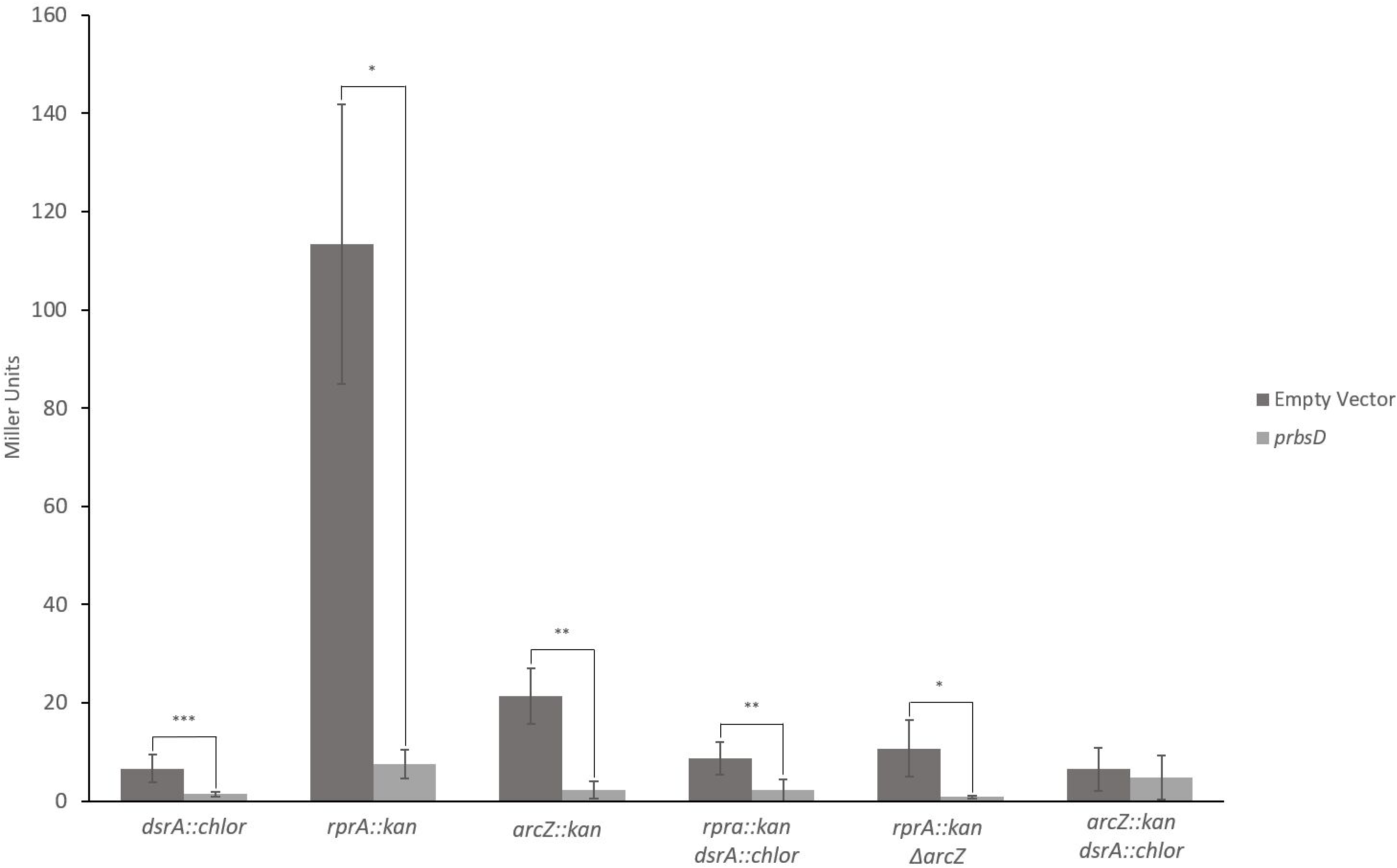
Both DsrA and ArcZ are necessary for RbsD regulation of RpoS. Beta-galactosidase assays of the RpoS750’-’LacZ fusion strain containing either single or double mutations in the sRNA genes of *arcZ, dsrA* and *rprA*. The presences of the p*rbsD* overexpression plasmid lowered the miller units in all the strains except for the double *dsrA arcZ* mutant (*, P < 0.05; **, P < 0.01; ***, P < 0.001). The single *dsrA* mutant diminished the response to p*rbsD* overexpression but did not abolish it completely.

We next performed experiments to determine if the binding site of DsrA on RbsD was necessary and sufficient to affect *rpoS* translation. The DsrA binding sites on RbsD has been identified (16) and we mutated six of the sixteen base pairs on RbsD that are involved in the binding. As seen in Figure 5a, levels of RpoS fusion activity were partially restored when these sites were disrupted. It was not possible to make the complementary mutations in DsrA since they would disrupt the interaction between DsrA and RpoS. Next, we cloned a small 80 base pair fragment of *rbsD* that contained the DsrA binding sites. This fragment lowered RpoS477’-‘LacZ levels to the same extent as the whole *rbsD* gene (Figure 5b). In sum, these data confirm that the mRNA of RbsD, in particular the small region binding DsrA, is responsible for RpoS regulation.

**Figure 5.**
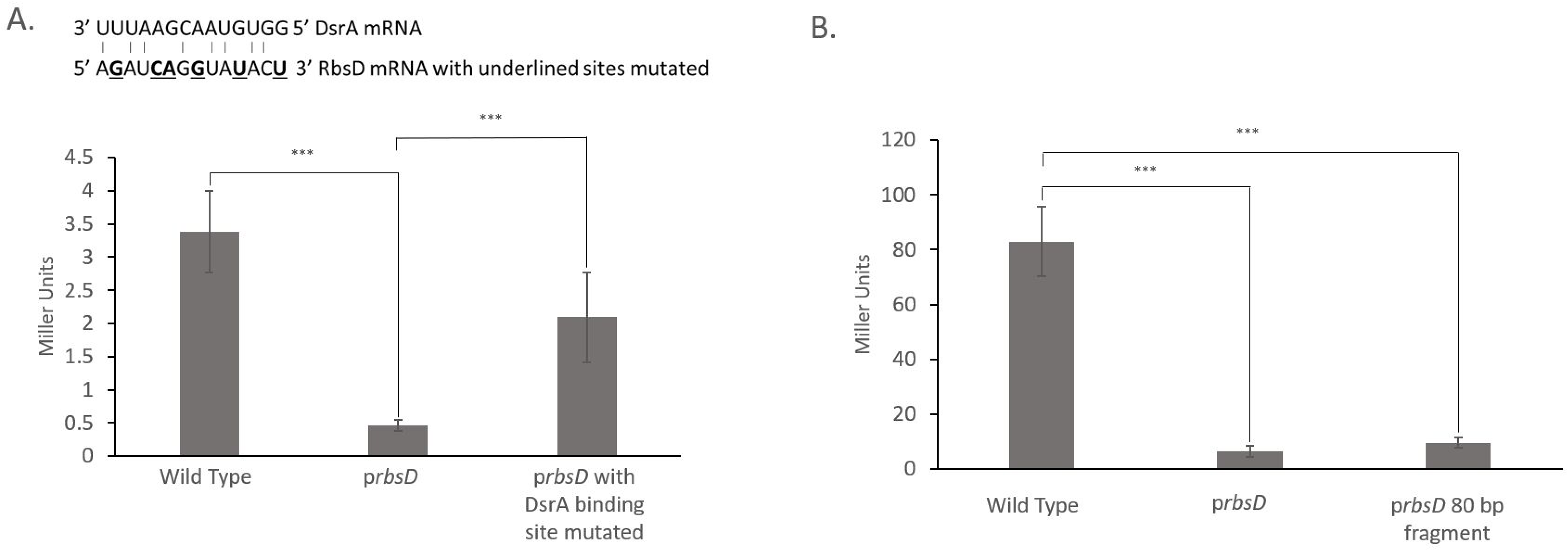
DsrA binding region on RbsD contributes to regulation. (A) Site-directed mutagenesis was used to create six silent point mutations on the 14-base sequence that has been shown to be the DsrA binding spot on RbsD. The bases underlined and in bold have been mutated in RbsD. Beta-galactosidase assays resulted in a *prsbsD* had lower fold effect on RpoS expression (***, P < 0.001). (B) An 80-base fragment of the *rbsD* gene with the native *rbsD* promoter was cloned into a plasmid, and overexpression of this fragment was shown to be sufficient for significantly lowering RpoS levels.

### Ribose lowers RpoS levels in a *rbsD* dependent manner

Since ribose activates RbsD expression, we next examined if ribose was a signal to lower RpoS levels (24). There was little effect of ribose in MC4100, which is expected since it has an *rbsR* mutation (*rbsR22)* which would render it insensitive to ribose (25). Subsequently, experiments were conducted in an MG1655 strain which has a wild type *rbsR*. Cells were grown exponentially in M63 glycerol and then transferred to either M63 glycerol or ribose for three hours. As seen in figure 6a, when ribose was added, the placRpoS’-‘LacZ fusion had three-fold lower activity. When *rbsD::kan* was introduced, there was no significant difference when ribose was added. In ribose conditions, the *rbsD::kan* allele had higher RpoS levels than wild type, however in LB conditions, there was no significant effect of *rbsD::kan* allele most likely since this operon is not highly expressed in rich media. Thus, ribose lowers RpoS levels through RbsD.

**Figure 6.**
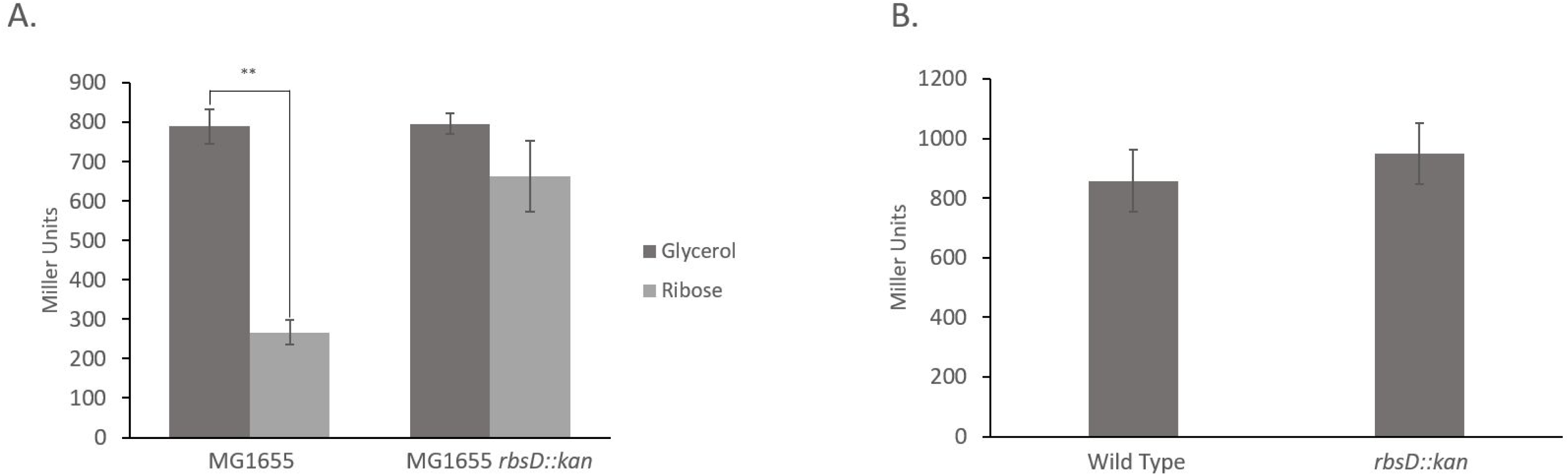
Ribose lowers RpoS levels in a manner that requires RbsD. MG1655 strains were grown exponentially in M63 glycerol and then transferred to either glycerol and ribose. Ribose lowered placRpoS’-LacZ levels in a statistically significant way in wild type MG1655 (**, P < 0.01). In the strain with *rbsD::kaņ* there was no significant effect of ribose on the placRpoS’-‘LacZ fusion.

## Discussion

Here we identify and describe a specific pathway for how ribose regulates RpoS. When the RbsD mRNA is expressed in ribose conditions, it lowers RpoS translation through a mechanism that requires the RpoS hairpin loop, the sRNA chaperone Hfq and the sRNAs DsrA and ArcZ. Interestingly, the RbsD pyranase protein is not required for this activity; it is sufficient to just express the part of the RbsD mRNA that binds DsrA to have the same effect as expressing the whole gene. Mutating six of the DsrA binding sites on RbsD partially abolished its effect on RpoS and it is possible that the residual effect is due to the other DsrA binding sites and/or due to ArcZ binding. The sugar ribose, which increases *rbsD* expression (24), lowered RpoS levels compared to similar growth in glycerol and this effect was dependent on RbsD. Ribose had little effect in MC4100, which has a RbsR mutation and thus constitutively expresses RbsD, but there was a strong effect in MG1655 which has a wild type RbsR. Thus, RbsR, which could be one of the polymorphisms that gives rise to the *E. coli* strain variation in RpoS levels and stress responses (26). Overall, our results delineating a clear ribose specific pathway for regulation of RpoS are in alignment with *rbsD* and *rbsR* mutations that have been identified in other screens as regulating RpoS (personal communication Susan Gottesman, Nadim Majdalani and Abbigale Perkins; (27)).

RpoS levels have long been known to respond to carbon conditions and this response often has been linked to central metabolism and ppGpp levels (4, 5). Here we have identified ribose as a major signal of nutrient availability, with its own specific pathway downregulating RpoS. The importance of ribose as a signaling molecule that triggers shifts in metabolism and stress response genes is becoming increasingly clear. Ribose is essential component of many of the pathways necessary for growth – including mRNA, rRNA, purines and pyrimidines, NADP/NAD - and can be used as a carbon source through the pentose phosphate pathway and glycolysis (28). Indeed, ribose is an essential component of ATP, which has also been suggested to be factor controlling RpoS degradation (5). Ribose availability correlates with purine regulation – when ribose is available, the purine salvage pathway is upregulated and in the absence of ribose, RbsR represses *de novo* synthesis of purines (29). Importantly, there are multiple regulatory RNA sponges embedded in the *rbs* operon (RbsD and RbsZ-S, described below), implying that transcription of this operon is crucial for signaling in several complex regulatory networks (30). Ribose is found in the enteric environment and specifically in the gut where Salmonella upregulates *rbsD* expression (31, 32). Finally, ribose can be released from the diet, the host and neighboring cells - especially in high levels from rRNA and mRNA but also from all the other cofactors, capsules and nucleic acid. *E. coli,* along with other symbionts and pathogens may have evolved to adapt quickly to take advantage of this carbon source (31, 33).

We have shown that RbsD plays a key role in the response to ribose as a dual function RNA; it produces both the ribose pyranase and also a regulatory RNA sponge that is separate of the protein (16). Indeed, the two roles are independent of each other since just an 80-base pair DsrA-binding region of RbsD is sufficient for its regulatory role. Similar to the model of competing endogenous RNAs (ceRNAs), we posit that when induced by ribose, the RbsD mRNA sequesters the sRNAs DsrA/ArcZ and Hfq so they are no longer available to open the *rpoS* hairpin loop for translation (34). There have numerous examples of bacterial RNAs acting as sponges (34–36). One of the first sponges described was the intergenic region of the *chbBC* transcript which bind to the chitobiose transport regulator sRNA ChiX. This interaction lowers the levels of ChiX (37, 38). While some bacterial sponges are co-degraded with their sRNAs, at this point it is not clear how the levels of DsrA RNA area affected by binding RbsD. The *chbBC* has a sequence that mimics ChiX target, ChiP and thus it acts like a decoy to titrate ChiX (37, 38). Likewise, on DsrA, the RbsD binding sequence overlaps with the RpoS binding sequence (16, 39), suggesting that there is direct competition between RbsD and RpoS for binding to DsrA (16, 39).

Post-translation regulation via a sponge mechanism can result in fast control of gene expression in response to rapidly changing nutrient conditions. There are several examples of RNA sponges connected to nutrient status (35, 36). In the ribose example, transcription of the *rbsDACB* and *rbsB*, leads to the 3’UTR of the transcript being processed into the RbsZ-S sponge RNA (another longer version of RbsZ is transcribed from a separate promoter). RbsZ-S binds to the sRNA RybB and stimulates its degradation so that it can no longer regulate the porins or RbsB itself. By acting as a sponge for RybB, RbsZ-S thus stimulates translation of RbsB in a positive autofeedback loop (30). Another sponge, the 3’UTR of *pspG*, downregulates the sRNA SpoT, which inhibits many catabolite repression genes. SpoT levels increase with glucose and decrease with cAMP levels, and it controls gene expression by inhibiting translation and lowers mRNA levels of its targets (40, 41). For amino acid transport, the SroC sponge titrates the sRNA GcvB, thus preventing it from decreasing expression of the amino acid transporters. There are a number of additional sRNAs that are either regulated by CRP/CAMP or themselves control genes involved carbon consumption (42, 43). Given the prevalence of sRNAs in regulation of carbon consumption, is possible that there are additional sponges also involved in fine tuning these sRNAs activity in response to carbon levels. New technologies are continuously being developing for identifying RNA sponges, which may lie in the 5 UTR, the coding region, the 3’ UTR, the intergenic or antisense regions of the genome (35, 36).

## Acknowledgements

We would like to thank the following people for helpful discussion: Susan Gottesman, Nadim Majdalani, Abbigale Perkins, Nicholas Wenner, Hannah Margalit, Liron Argaman, Melanie Berkmen and Thomas Silhavy.

## Materials and methods

### Bacterial strains and culture conditions

Bacterial strains used in this study are derived from *E. coli* K-12 strains MC4100 and MG1655. All the strains and plasmids are listed in Table S1. All strains were grown in Luria Broth or M63 salts at 37 °C. The M63 media was supplemented with 0.001% vitamin thiamine, 0.0001% biotin and either glucose, glycerol or ribose at 0.4%. For growth in LB, the following concentrations of antibiotics were used: 100 μg/ml ampicillin, 10 μg/ml chloramphenicol and 50 μg/ml kanamycin.

The strains with *rprA::kan*, *dsrA::chlor* and *rbsD::kan* strains were created using P1*vir* phage transductions as previously described by Miller (44). The *<u>arcZ</u>*::*kan* and *arcZ* deletion were created with the lamda red recombination system (45). The primers contained 60 bases pairs of homologies for the downstream and upstream region of the arcZ gene as well as homology with the kanamycin resistance gene (ArcZkanfor and ArcZkan) PCR was carried out with the Q5 2X Master Mix (New England Biolabs). The mixture was run in the thermocycler with a denaturation temperature of 95°C, annealing temperature of 55°C, and an extension temperature of 72°C for 25 cycles. The PCR cassette was transformed into the strain carrying pSIJ8. To test kanamycin resistant colonies for successful recombination, *arcZconfirm* forward and reverse primers were used on gDNA extracted from overnight cultures and the products were run on agarose gel electrophoresis and sequenced (Azenta). The kanamycin cassette was removed through the FRT sites as previously described (45) and the region was amplified and sequenced for confirmation of the recombination.

Plasmids were also cloned to include the *rbsD* promoter with either the full length *rbsD* gene or with only the 80 base pair region surrounding the DsrA binding sites onto using TOPO TA cloning. The regions were amplified with PCR with Taq Polymerase and cloned according to the pCR™8/GW/TOPO™ TA protocol (Thermo Fisher Scientific).

The site directed mutagenesis of the *rbsD* gene to introduce the stop codon and the mutated *dsrA* binding sites was carried out using the Q5 mutagenesis kit (New England Biolabs) and selection on kanamycin plates. The primers for mutagenesis of the stop codon were RbsDstopfor and RbsDstoprev while the DsrA binding site mutagenesis primers were RbsDmutfor and RbsDmutrev. These primers changed made 6 silent mutations that affected the DsrA binding sites. All constructs were verified with Sanger sequencing.

### Overexpression library and screen

The overexpression library was created by isolating genomic DNA (Qiagen) and cutting it with EcoRV. Fragments from 500 bp to 1000 bp were excised from an agarose gel and cloned into the pSMART LCKan plasmid (Lucigen) by transforming into competent cells. Colonies from the cloning were grown overnight at 37 °C scrapped off plates and miniprepped to create the library. For the screen, this library was transformed by electroporation into CP119 which carried the RpoS750’-‘lacZ reporter strain and selected on X gal kanamycin plates after overnight growth at 37 °C. Light blue candidate colonies were streaked before miniprepping and re-transforming into the original RpoS750’-‘lacZ reporter strain.

### Betagalactosidase assays

*lacZ* expression in the experiments with all the different reporters, was determined by β-galactosidase activity assay as described by Miller (44). For LB growth assays, bacterial cultures were grown overnight, diluted 1:200 and the 1.5 mls was collected for the assay between OD_600_ = 0.3 to 0.4. At least three biological triplicates were done for each assay and data were plotted as means with standard errors (SEMs). For the M63 experiments, cells were grown in M63 0.2% glycerol overnight and then diluted 1:10 in fresh M63 glycerol and grown shaking for 2 hours at 37°C. Cells were spun down at 6000 rpm and then switched to M63 0.4% ribose or glycerol and grown shaking for 2 hours at 37°C. 1.5 mls of each culture was collected for the assay.

### qRT-PCR

To measure the effect of *prbsD* on RpoS mRNA, cultures were grown overnight in LB with antibiotics, diluted in LB 1:200 and then samples were collected between OD_600_ = 0.3 to 0.4. RNA was extracted (Qiagen), treated with DNAase (New England Biolabs). cDNA was created with SuperscriptII (Thermo Fisher Scientific) and random hexamers. qRT_PCR was carried out with SYBER Green qPCR Master Mix (Thermo Fisher Scientific) with primers previously described (46). At least three biological triplicates were done for each assay and triplicate technical assays were carried out.

### Western blots

Overnight cultures were diluted 200-fold into fresh LB media and incubated and incubated at 37°C with shaking at 225 rpm. A 1.5 ml-sample was collected from each culture when the OD_600_ reached between 0.3 and 0.4 and resuspended in sodium dodecyl sulfate (SDS) loading buffer in an OD_600_/10 volume. After electrophoresis, proteins were transferred onto nitrocellulose membranes and probed with a 1:2,000 dilution of anti-RpoS antibody (Biolegend). For secondary antibody, rabbit anti-mouse horseradish peroxidase was used 1:1000. (Abcam).

